# Massive contractions of Myotonic Dystrophy Type 2-associated CCTG tetranucleotide repeats occur via double strand break repair with distinct requirements for helicases

**DOI:** 10.1101/2023.07.06.548036

**Authors:** David Papp, Luis A. Hernandez, Theresa A. Mai, Terrance J. Haanen, Meghan A. O’Donnell, Ariel T. Duran, Sophia M. Hernandez, Jenni E. Narvanto, Berenice Arguello, Marvin O. Onwukwe, Kara Kolar, Sergei M. Mirkin, Jane C. Kim

## Abstract

Myotonic Dystrophy Type 2 (DM2) is a genetic disease caused by expanded CCTG DNA repeats in the first intron of *CNBP*. The number of CCTG repeats in DM2 patients ranges from 75-11,000, yet little is known about the molecular mechanisms responsible for repeat expansions or contractions. We developed an experimental system in *Saccharomyces cerevisiae* that enables selection of large-scale contractions of (CCTG)_100_ within the intron of a reporter gene and subsequent genetic analysis. Contractions exceeded 80 repeat units, causing the final repetitive tract to be well below the threshold for disease. We found that Rad51 and Rad52 are required for these massive contractions, indicating a mechanism that involves homologous recombination. Srs2 helicase was shown previously to stabilize CTG, CAG, and CGG repeats. Loss of Srs2 did not significantly affect CCTG contraction rates in unperturbed conditions. In contrast, loss of the RecQ helicase Sgs1 resulted in a 6-fold decrease in contraction rate with specific evidence that helicase activity is required for large-scale contractions. Using a genetic assay to evaluate chromosome arm loss, we determined that CCTG and reverse complementary CAGG repeats elevate the rate of chromosomal fragility compared to a low-repeat control. Overall, our results demonstrate that the genetic control of CCTG repeat contractions is notably distinct among disease-causing microsatellite repeat sequences.

## Introduction

Repeat expansion diseases are caused by excessively long tracts of simple DNA sequence repeats, also called microsatellites, within specific genes. Two types of myotonic dystrophy (DM) are examples of such diseases (*1*). DM1 is caused by long (>50) CTG trinucleotide repeats (TNRs) in the 3’ UTR of *DMPK*, whereas DM2 is caused by long (>75) tetranucleotide CCTG repeats in the first intron of *CNBP*. Transcription of these expanded alleles results in cellular toxicity through RNA gain-of-function effects (*2*). Repeat associated non-ATG (RAN) translation products were observed in brain tissues of DM1 and DM2 patients (*3, 4*), suggesting that aberrant proteins may also contribute to pathophysiology. Disease symptoms include inability to relax muscles after they have contracted, muscle weakness, and cardiac conduction problems. However, no cure or specific therapy is currently available for either disease, with treatment relying on symptom management.

Individuals with DM1 or DM2 may have thousands of tandem repeats within the affected gene. Through studies using model organisms and cell culture systems, there have been substantial advances in understanding CTG and reverse complementary CAG instability (expansions and contractions) (*5–8*). CAG/CTG repeats form stable hairpins and slipped strands *in vitro*, and formation of these secondary structures *in vivo* are proposed to instigate various molecular mechanisms of instability (*9–11*). For example, polymerase slippage followed by hairpin formation on the nascent strand during DNA replication or repair synthesis would lead to an expansion. In contrast, structure formation on the template strand, which is then bypassed during DNA synthesis, would result in repeat contractions. Additionally, CAG/CTG repeats cause replication fork stalling (*12, 13*) and elevated DNA double strand break (DSB) formation (*14, 15*), and their recovery or repair via homologous recombination could lead to expansions or contractions depending on the fidelity of steps such as strand annealing (*16*).

Understanding the molecular mechanisms of contractions is important to evaluate whether manipulating this process would be a viable approach to treat repeat expansion diseases (*17*). Strategies to induce targeted repeat contractions using either small molecules that specifically bind to slipped strand intermediates (*18*) or sequence-specific endonucleases to generate DSBs at the repeat locus (*19*) were successful in experimental systems with CAG/CTG repeats. It is unclear whether these strategies would apply generally to all disease-causing DNA repeat sequences or whether the underlying mechanisms of large-scale contractions for different repeats are also distinct. Specifically, we designate the descriptors massive or large-scale as the number of repeats that would result in a decrease from symptomatic to unaffected lengths (*i.e.* a change in >45 repeats for DM2 as opposed to loss of just a few repeats).

In contrast to CAG/CTG repeats, much less is known about the mechanisms of CCTG and reverse complementary CAGG repeat instability. This is a pressing question since individuals with DM2 can have 75-11,000 CCTG repeats in *CNBP* with an estimated average of 5000 repeats (*20*). DM2 is also the only repeat expansion disease currently known to be caused by tetranucleotide repeats. *In vitro* studies with chemical and enzymatic probing showed that CAGG repeats form stable hairpin structures with specific base-pairing propensity, which was not prominent for the CCTG orientation (*21*). CAGG/CCTG repeats were also shown to form slipped-strand structures through denaturation and reduplexing experiments (*22*). In addition, NMR analysis indicated that CCTG repeats form hairpin and dumbbell structures that are much more fluid and dynamic (*23*), displaying an ability to change between different conformations and shift along the DNA duplex. This fluidity is proposed to increase the likelihood that CCTG repeats could escape DNA repair (*24, 25*).

Previous studies investigating CAGG/CCTG repeat instability *in vivo* have relied on plasmid-based systems using *E. coli* and COS-7 mammalian cell culture (*21, 26*). The frequency of expansions and contractions increased with longer starting lengths, and the orientation with CAGG repeats on the leading strand template showed greater instability. Another plasmid study using *E. coli* found that instability was not dependent on repeat orientation (*27*). In the current study, we developed a yeast experimental system to investigate instability of CAGG/CCTG repeats within a chromosomal context. This system offers an additional advantage of enabling the selection of repeat contractions that exceed 80 repeats in length, for example from (CCTG)_100_ to (CCTG)_20_. Through genetic analysis, we found that homologous recombination is required for large-scale CCTG repeat contractions. We also demonstrate that CAGG/CCTG repeats elevate the rate of double strand break formation using a genetic assay for chromosomal arm loss. Altogether, genes that have been well-characterized to control CAG/CTG repeat instability do not affect CAGG/CCTG repeats in the same manner, indicating that the precise genetic control of DNA repeat instability is unique to the specific microsatellite sequence.

## Materials and Methods

Long CCTG repeats were cloned using CCTG and CAGG oligonucleotides designed with 5’ non-palindromic restriction sites, following a previously described strategy to clone TNRs (Supplementary Fig. 1) (*28, 29*). To construct the *URA3* reporter gene, CCTG repeats were cloned into a plasmid described as (GAA)_0_ (*30*) that had been cut with NaeI. The (CCTG)_100_ plasmid is designated p18 and contains an overall intron length of 1047 bp.

Additional information regarding plasmid cloning, yeast strain construction, spot assays, fluctuation analysis, and RT-qPCR can be found in Supplemental Information.

## Results

### A System to Investigate CAGG/CCTG Repeat Contractions

To investigate CAGG/CCTG repeat instability, we employed a system previously used to study large-scale GAA repeat expansions (*31*) and contractions (*32*), which are responsible for Friedreich’s Ataxia (FA). The yeast system mimics the intron location of GAA repeats in the gene responsible for FA, wherein *URA3* under the control of its native promoter is artificially split by an intron containing (GAA)_100_ repeats and used for forward selection. Large-scale GAA expansions increase the intron length beyond the yeast splicing threshold of ∼1 kb, which impairs *URA3* splicing and renders cells resistant to the drug 5-fluoroorotic acid (5-FOA). Cells can also become 5-FOA^R^ through spontaneous point mutations, insertions and deletions, or other mechanisms that inactivate *URA3* expression. We adapted the system by cloning (CCTG)_100_ repeats into the artificial intron of *URA3* (Fig. 1a). The cassette was integrated into chromosome III, ∼1 kb away from *ARS306*, a replication origin that is efficient and activated early in S-phase. In these cells, CCTG is on the sense strand, as it is for *CNBP*, and lagging strand template for DNA replication. We also constructed an equivalent yeast strain with (CAGG)_100_ on the lagging strand template to investigate the effect of repeat orientation.

**Figure 1.**
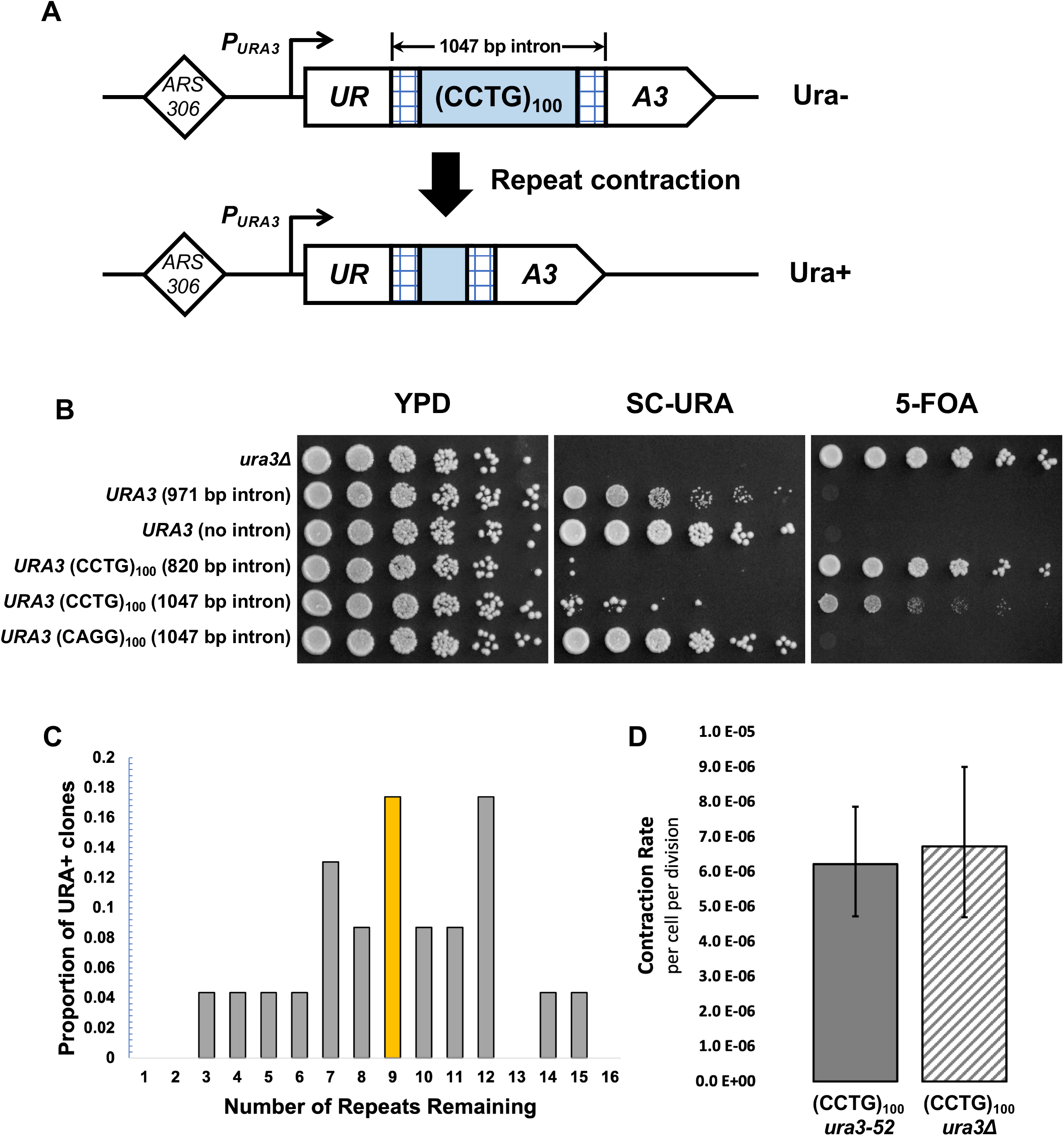
Experimental system to study large-scale CCTG repeat contractions *in vivo*. (A). DNA repeats are cloned into the artificial intron, derived from *ACT1*, of a *URA3* reporter gene. The reporter gene is integrated ∼1 kb downstream of the replication origin *ARS306*. The starting strain with (CCTG)_100_ is Ura-. Repeat contraction renders the cells Ura+. (B) Spot assays to evaluate the growth phenotypes of yeast strains with (CCTG)_100_ and (CAGG)_100_ repeats as well as non-repetitive DNA. (C) Distribution of remaining repeat length in Ura+ clones, evaluated by PCR and Sanger sequencing. The median number of repeats was nine. (D) Rate of large-scale contraction for (CCTG)_100_ strains, shown with 95% confidence intervals. Rate is calculated using the number of Ura+ clones beginning with 12 independent cultures and the Ma–Sandri–Sarkar maximum-likelihood estimator with a correction for sampling and plating efficiency.

Unexpectedly, we found that the two orientations led to distinct growth phenotypes with respect to uracil auxotrophy (Fig. 1b), despite the total intron length of both strains being identical (1047 bp). The (CAGG)_100_ orientation resulted in a Ura+ phenotype equivalent to *URA3* with no intron: growth occurred on synthetic media lacking uracil whereas no growth occurred on 5-FOA media. In contrast, the (CCTG)_100_ orientation showed a predominantly Ura-phenotype with some distinct Ura+ papillae. To exclude the possibility that the (CCTG)_100_ strain was Ura-because the intron length exceeded the yeast splicing threshold, we generated yeast with (CCTG)_100_ but a shorter overall intron length (820 bp). The growth phenotype was also Ura-in this modified strain (Fig. 1b). We used the (CCTG)_100_ strain with the 1047 bp intron length for all subsequent analyses of the CCTG orientation.

We hypothesized that the (CCTG)_100_ orientation inhibited functional *URA3* expression and that Ura+ clones in the spot assay were due to spontaneous repeat contractions that restored its expression. We performed RT-qPCR to evaluate how transcript levels corresponded to uracil auxotrophy. We observed that a strain with no intron showed >100-fold increase in *URA3* transcript abundance compared to the no-repeat intron (971 bp) and (CCTG)_100_ strains (Fig. 2). The (CCTG)_100_ strain showed a decrease (2.6-fold) in spliced *URA3* transcript compared to the no-repeat intron strain, providing evidence that a splicing defect in the CCTG orientation contributes to its Ura-phenotype. We also analyzed gene expression in strains with (CCTG)_22_, (CCTG)_43_, (CCTG)_78_ in the *URA3* intron and found an inverse relationship between repeat length and abundance of the spliced *URA3* transcript (Supplementary Fig. 2), providing further evidence that shorter CCTG repeat tracts restore functional *URA3* gene expression in this system. In contrast, the (CAGG)_100_ strain showed a 3.8-fold increase in spliced *URA3* transcript compared to the no-repeat intron strain.

**Figure 2.**
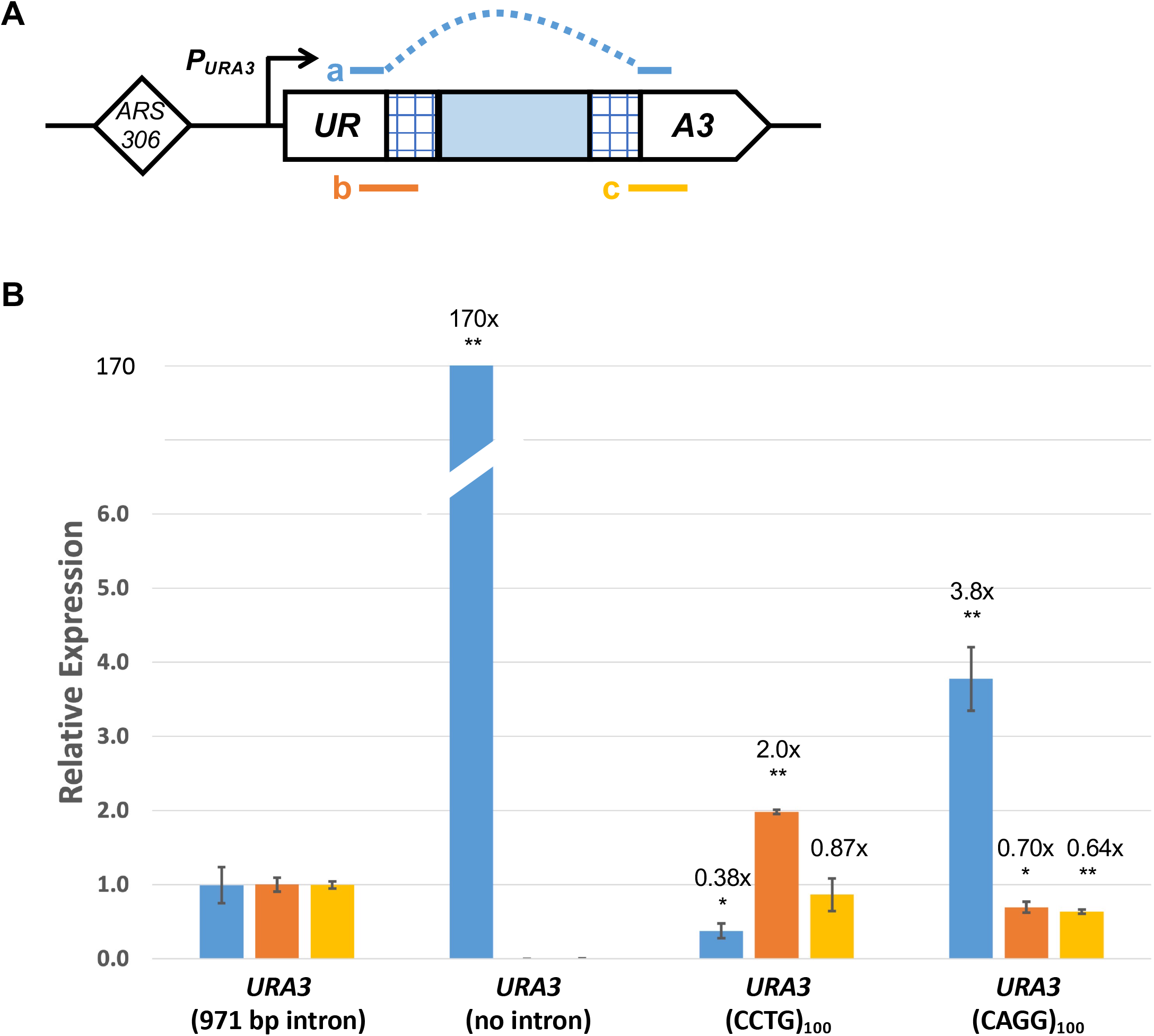
Comparison of *URA3* reporter gene expression in strains with and without CAGG/CCTG repeats. (A) Representation of PCR amplicons to evaluate: (a) spliced *URA3*, (b) 5’ unspliced *URA3*, and (c) 3’ unspliced *URA3*. Dashed line denotes that primers will not anneal to the intron sequence. (B) Relative expression of spliced and unspliced *URA3* transcripts compared to *ACT1* endogenous control in four yeast strains. Bar graph colors correspond to amplicon colors as in (A). The relative expression in the long intron strain was set to 1 for comparison, and numerical fold-difference is in comparison to the long intron strain. At least three biological replicates of cDNA for each strain were used for the analysis. The mean relative expression values and standard error are plotted. Statistical significance was evaluated by unpaired *t* test (* *p*<0.05, ** *p*<0.01)

To measure the rate of Ura+ clone formation, we conducted fluctuation tests on the (CCTG)_100_ orientation strain. After performing PCR analysis on 24 Ura+ clones using primers flanking the repeats and sequencing the PCR products, we found that the remaining number of repeats ranged from 3-15, with a median of 9 repeats (Fig. 1c). This indicates a massive net contraction of over 80 repeats. We calculated the rate of Ura+ clone formation to be 6 x 10^-6^ per replication (Fig. 1d), which we designate as the wild type (WT) rate of large-scale contraction based on the PCR and sequencing analyses. To confirm that Ura+ clones were not due to gene conversion events with the endogenous *ura3-52* allele on chromosome V, which contains a Ty1 retrotransposon insertion, we integrated the (CCTG)_100_ cassette into an isogenic background with a full *URA3* deletion. We observed an equivalent rate of Ura+ clone formation (Fig. 1d), indicating that they are primarily (if not exclusively) due to large-scale contractions. Next, we focused on delineating the genetic control of large-scale (CCTG)_100_ contractions.

### Impairment of lagging strand synthesis does not affect large-scale CCTG repeat contractions

The previous study investigating large-scale contractions of GAA repeats found that contractions occur during DNA replication rather than DNA repair pathways (*32*). The median contraction size was ∼60 repeat units, and a mechanism involving bypass of a transient triplex DNA structure during lagging strand synthesis was proposed. Notably, there was a >40-fold increase in large-scale GAA contractions following loss of Rad27, a 5ʹ to 3ʹ exonuclease and 5ʹ flap endonuclease that processes single-stranded flaps during Okazaki fragment maturation. In contrast, we found that large-scale CCTG contraction rate in the *rad27Δ* strain was indistinguishable from WT (Fig. 3, Supplemental Table 1). Furthermore, we found only a small increase (1.6-fold) by knocking out the processivity unit of DNA polymerase δ (*pol32Δ*), which was not significant. Because these key proteins in DNA replication and lagging strand DNA synthesis showed no effect on large-scale CCTG contractions, we focused our subsequent genetic analysis on DNA repair pathways.

**Figure 3.**
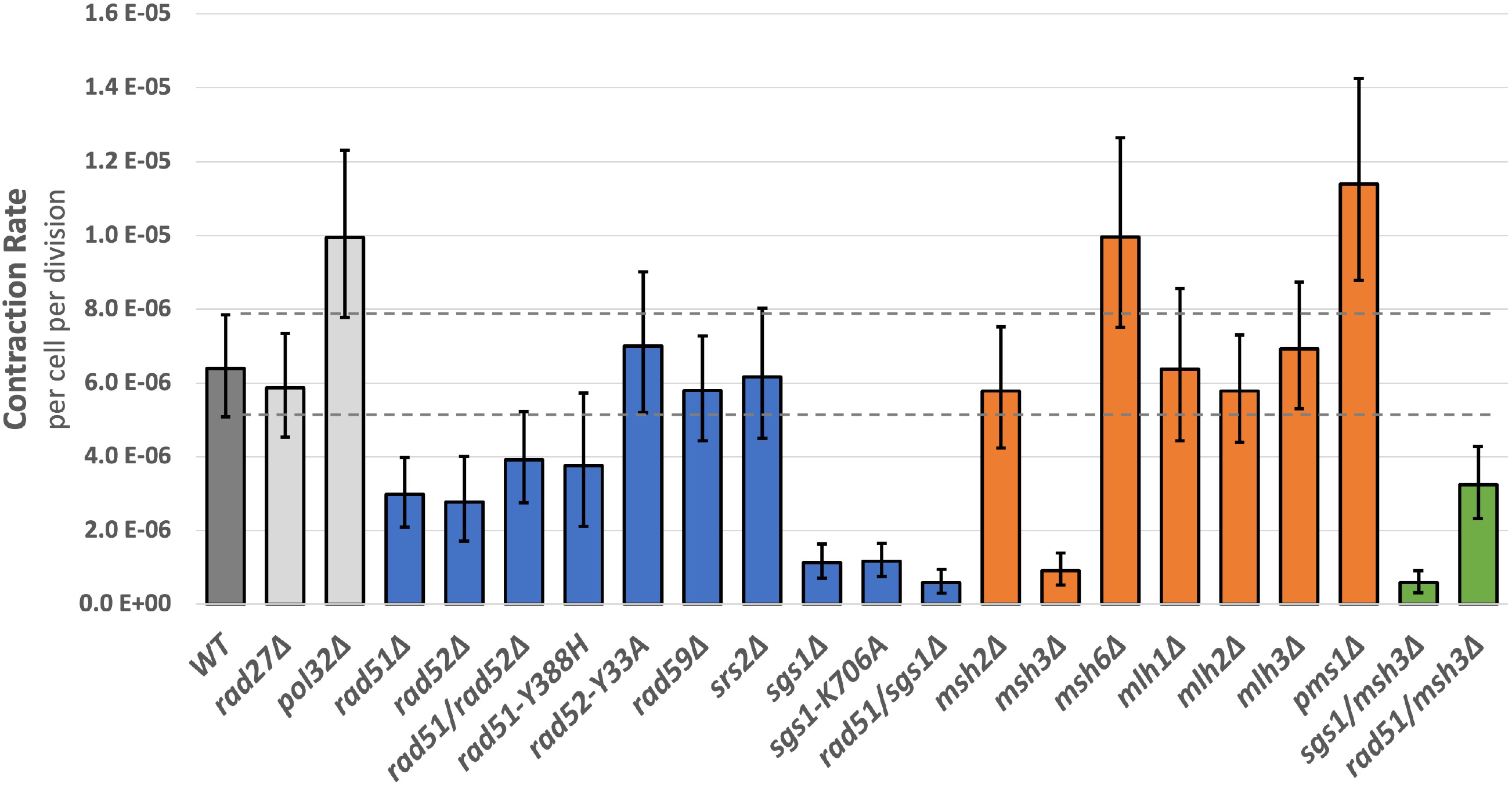
Genetic analysis of large-scale CCTG repeat contractions. Rate of large-scale contraction for (CCTG)_100_ strains, shown with 95% confidence intervals (dashed lines for WT strain). Rate is calculated using the number of Ura+ clones beginning with 12 independent cultures and the Ma–Sandri–Sarkar maximum-likelihood estimator with a correction for sampling and plating efficiency. Two rates are considered statistically different when the 95% confidence intervals do not overlap.

### Homologous recombination is required for large-scale CCTG repeat contractions

Various DNA repeats have been shown to elevate homologous recombination (HR) in plasmid-based systems, resulting in expansions and contractions. For example, using an *E. coli* intramolecular plasmid system, CAGG/CCTG repeats increased recombination crossover frequencies with expansions being more prevalent than contractions (*26*). Similar results were observed for CAG/CTG (*33*) and GAA/TTC (*34*) repeats using this *E. coli* system, though contractions were more prevalent. The role of HR on chromosomal DNA repeats is more varied depending on tract length and microsatellite sequence. Loss of Rad51 recombinase and Rad52, which facilitates Rad51 loading, had no effect on short tracts of (CAG)_13_ or (CAG)_25_ repeats (*35, 36*) whereas it did enhance small-scale expansions (*37*) and contractions (*38*) of (CAG)_70_, indicating a protective effect of HR. In contrast, large-scale expansions of (CAG)_140_ were dependent on HR through a proposed break-induced replication (BIR) mechanism (*39*). However, loss of Rad52 had no effect on large-scale GAA repeat expansions (*31*) and a mildly protective effect (2-fold increase in *rad52Δ*) on large-scale GAA contractions (*32*).

We observed a 2.2-fold decrease in large-scale contraction rates in *rad51Δ* and 2.3-fold decrease in *rad52Δ* compared to WT (Fig. 3), demonstrating that HR promotes this process. We found that this decrease was consistent using two analytical methods: non-overlapping 95% confidence intervals, which is standard for this analysis, and comparison of multiple trials encompassing 59 independent cultures (Supplementary Fig. 3). Furthermore, we tested whether Rad51 and Rad52 might have a compensatory effect or overlapping functions. The *rad51Δ rad52Δ* double mutant showed a comparable, epistatic (and not synergistic) 1.6-fold decrease to the single mutants, indicating that their roles are in the same genetic pathway. We tested specific mutants to investigate the roles of these proteins further. Rad51-Y388H is defective in interactions with Rad52 as well as Srs2 helicase (*40*). This mutant demonstrated a 1.7-fold decrease in large-scale contraction rate. Rad52-Y33A (see Supplemental Methods for nomenclature) is defective in DSB repair, as demonstrated by sensitivity to γ-irradiation, but proficient in spontaneous mitotic recombination such as heteroallelic recombination. The large-scale contraction rate of Rad52-Y33A was indistinguishable from WT. We further evaluated the role of the Rad52 homolog Rad59, which is required for single strand annealing, and found no significant difference in large-scale contraction rate compared to WT.

### Srs2 helicase is not required for large-scale CCTG repeat contractions in unchallenged cells

Srs2 is a DNA helicase that functions in many aspects of DNA replication, repair, and recombination. Notably for DNA repeat instability, Srs2 can unwind CAG, CTG, and CGG substrates *in vitro* (*41*). Two-dimensional gel-electrophoretic analysis of replication intermediates demonstrated that Srs2 is required *in vivo* for replication fork progression through these TNRs (*42*). Loss of Srs2 function caused an increase in small-scale expansions of (CAG)_25_, (CTG)_25_, and (CGG)_25_ repeats (*36*) and non-selective expansions and contractions of (CTG)_55_ (*42*), though it had no effect on large-scale expansions of (CAG)_140_ (*39*). Because CCTG/CAGG repeats can also form hairpin structures, we investigated loss of Srs2 function in our assay, predicting that there would be an increase in large-scale contractions if Srs2 unwound CCTG/CAGG repeats due to replication bypass of structures formed on the template strand. We found that the rate of large-scale CCTG contractions in *srs2Δ* did not differ from WT (Fig. 3).

Because Srs2 has an anti-recombinase role and displaces Rad51 filaments during HR, we tested whether loss of Srs2 function would influence large-scale CCTG contractions when cells were challenged with replication stress, specifically DSBs. When *srs2Δ* cells were treated with camptothecin (CPT), a topoisomerase I inhibitor that results in breaks during DNA replication, rates of large-scale CCTG contractions increased over a range of concentrations (Supplementary Fig. 4). For example, comparing *srs2Δ* to WT, there was a 4.8-fold increase at 10 μM CPT and 5.7-fold increase at 50 μM CPT. In addition, 50 μM CPT increased the contraction rate of *srs2Δ* cells 11.8-fold compared to untreated *srs2Δ* cells, revealing a dose-dependent effect of DNA damage on the role of Srs2 function in large-scale CCTG contractions. Furthermore, we observed a similar increase in contraction rates with hydroxyurea (HU) treatment (Supplementary Fig. 4), which causes replication fork stalling and uncoupling of the replication fork polymerases and helicase that can result in DSBs. At 50 mM HU, *srs2Δ* showed a 3.1-fold increase compared to WT. For both CPT and HU treatments, there was also a ∼2-fold increase in contraction rate at all doses in the WT strain compared to no treatment. We propose that the increase is indicative of DSBs that are instigated during S phase and aberrantly repaired to generate large repeat contractions (see Discussion).

### Sgs1 helicase is required for large-scale CCTG repeat contractions

The RecQ helicase, Sgs1, plays various roles in DNA repair and recombination in budding yeast (*43*). During recombination, Sgs1 plays an early role in long-range resection at a DNA double strand break and was shown to be required for heteroduplex rejection during single strand annealing (*44*). Through biochemical assays, purified Sgs1 was demonstrated to unwind a broad range of DNA structures including Holliday junctions (*45*) and G-quadruplexes (*46*). Long (CTG)_75_ repeats were shown to be stabilized by Sgs1, as loss of function led to increased expansions and contractions under non-selective conditions (*42*).

The absence of Sgs1 had a considerable effect on large-scale CCTG contraction rate, as *sgs1Δ* showed a 5.7-fold decrease compared to WT (Fig. 3). Interestingly, Sgs1 is required for large-scale contractions even under unperturbed conditions, which is opposite the effect displayed by Srs2 when treated with replicative stressors. To better understand the role of Sgs1 in repeat contractions, we constructed and analyzed a helicase-defective mutation Sgs1-K706A. This strain also displayed a 5.5-fold decrease in contraction rate compared to WT (Fig. 3), indicating that Sgs1 helicase activity is required for its effect on large-scale CCTG contractions. We evaluated contraction rate in the *rad51Δ sgs1Δ* double mutant and found a 10.8-fold decrease in contraction rate compared to WT (Fig. 3). This was a slight decrease compared to the *sgs1Δ* mutant, though the 95% confidence intervals are overlapping.

Having established a requirement for Sgs1 on large-scale CCTG repeat contractions, we investigated whether its role was through facilitating 5’ DNA end resection, a key processing step to promote double strand break repair by homologous recombination. The Mre11-Rad50-Xrs2 (MRX) complex localizes to a DSB, upon which the Mre11 nuclease can initiate resection. Sae2 has also been shown to be involved in short-range resection, though both Mre11 and Sae2 have been shown to be dispensable for resection of DNA ends overall. Long-range, more processive resection is mediated by Sgs1-Dna2 or Exo1 pathways, which may have redundant roles. We found that single mutants of *mre11Δ*, *sae2Δ*, and *exo1Δ* each displayed large-scale contraction rates that were not significantly different from WT (Supplementary Fig. 5). Sgs1 interacts with Top3 and Rmi1 to form the STR complex, which enhances DNA end resection and resolves recombination intermediates. We did not observe an effect on large-scale contraction rates in the *rmi1Δ* mutant.

Because the nuclease domain of Dna2 is essential, its role in DNA resection could not be evaluated through direct knockout. We tested the Dna2-H547A mutant, which is described as nuclease-attenuated, though it maintains ATPase and helicase activity (*47*). We found that the mutant displayed a 12-fold elevated rate, suggesting that Dna2 nuclease activity plays a protective role in CCTG repeat contractions (Supplementary Fig. 5).

### Mismatch repair proteins are differentially required for large-scale CCTG repeat contractions

DNA mismatch repair (MMR) is initiated by the MutSα and MutSβ complexes, formed by the Msh2-Msh6 and Msh2-Msh3 heterodimers, respectively. MutSα recognizes single-base mismatches, while MutSβ recognizes insertion/deletion loops. MMR proteins have been implicated in CAG/CTG repeat instability. Specifically, CAG/CTG trinucleotide repeats were stabilized in the *msh3Δ* mutant (*48*). Loss of Msh2 and Msh3, though not Msh6, were shown to decrease expansions in mouse models (*49–51*). These results were corroborated in a human cell culture system (*52*). However, loss of MMR proteins had no effect on large-scale (>60 repeats) expansions of CAG/CTG repeat in a yeast experimental system (*39*).

In evaluating large-scale CCTG contractions, we found that loss of MMR proteins demonstrated differential effects (Fig. 3). The strongest effect was displayed by *msh3Δ*, which showed a 7.0-fold decrease in contraction rate. In contrast, *msh2Δ* showed no difference and *msh6Δ* a slight increase (1.6-fold) in contraction rate, both of which were not significant. Following mismatch recognition, proteins in the MutL family function as heterodimers to catalyze repair. They also have roles outside of MMR such as in meiotic recombination (*53*). Large-scale contraction rates in *mlh1Δ*, *mlh2Δ*, *mlh3Δ*, and *pms1Δ* single mutants were not significantly different from WT.

To evaluate the relationship of Msh3 with Sgs1 and Rad51, we constructed double mutants and tested their effect on large-scale CCTG repeat contractions. The *sgs1Δ msh3Δ* mutant showed a 11-fold decrease in contraction rate compared to WT, a slight decrease compared to the single mutants though the 95% confidence intervals overlap. In contrast, the *rad51Δ msh3Δ* mutant showed a rate (2.0-fold decrease) comparable to the *rad51Δ* single mutant, evidence of an epistatic relationship where Rad51 functions upstream of Msh3 in the genetic pathway. We synthesize these genetic results in the Discussion.

### CCTG repeats increase chromosomal fragility in an orientation-dependent manner

To confirm DSB formation caused by CAGG/CCTG repeats, we used a genetic assay (*54*) to determine the rates of chromosomal fragility in both repeat orientations. This assay works by integrating DNA repeats into a relocated *lys2* locus on ChrV (Fig. 4a), where DNA breakage can lead to a telomere-proximal deletion (arm loss). This region does not contain essential genes but does contain *CAN1* and *ADE2*. This allows for selection of canavanine resistant (Can^R^) and Ade-clones, which appear red, and permits subsequent calculation of DNA fragility rates. For this arm loss assay, we designate repeat orientation as previously described and name the repeat sequence that is on the lagging strand template for DNA replication. This orientation is determined by *ARS507*, which was characterized by 2D gel analysis of replication intermediates (*55*).

**Figure 4.**
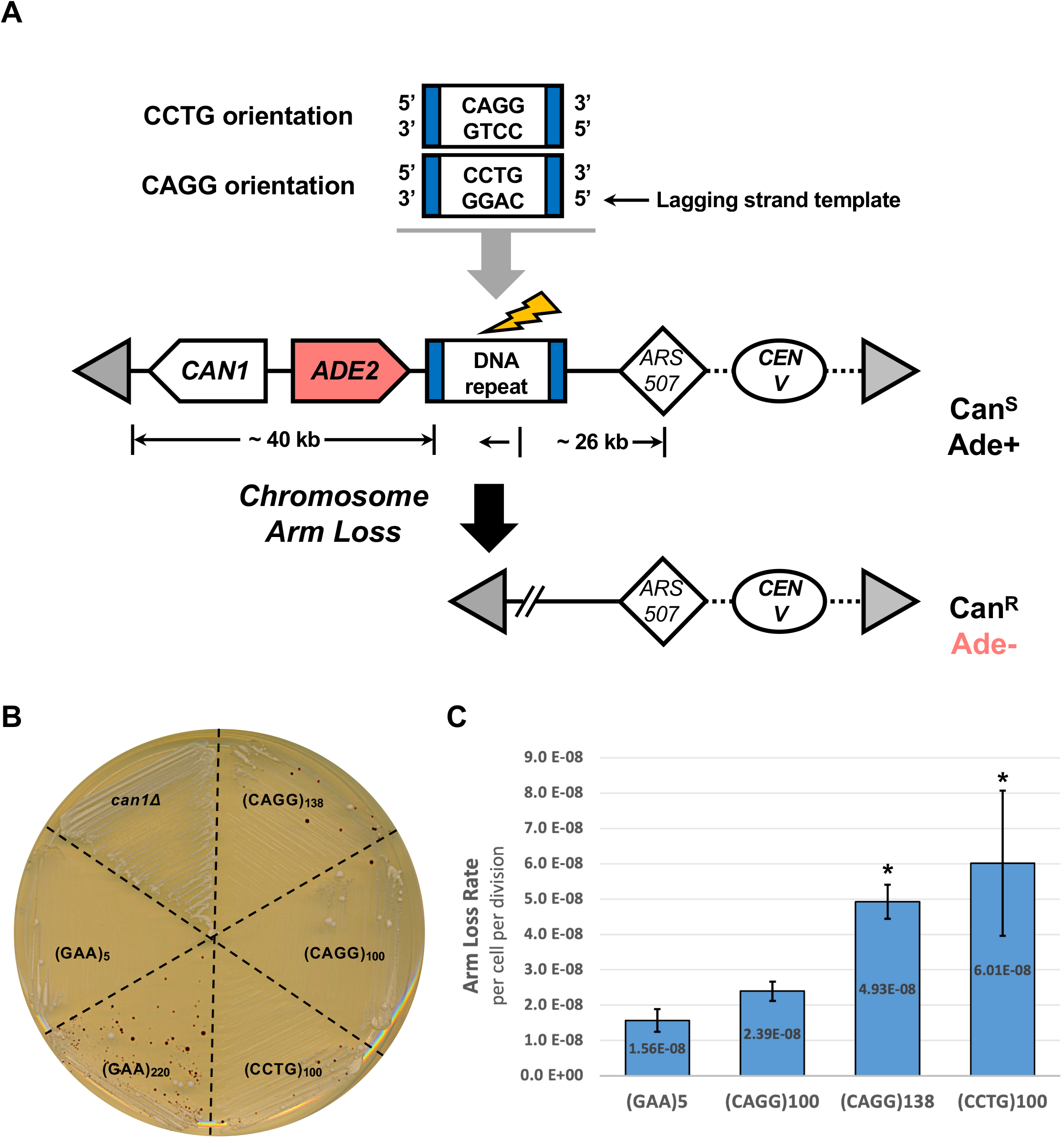
CCTG repeats elevate chromosomal fragility in an orientation-dependent manner. (A) Experimental system to study repeat-induced DNA fragility via chromosomal arm loss. CCTG and CAGG repeats were integrated at the reporter locus, *lys2* (blue). Double strand break formation followed by telomere addition (gray triangle) will result in increased arm loss frequency of the left chromosomal arm. This region (∼40 kb) contains no essential genes and the selectable markers *CAN1* and *ADE2*. Mutations in these genes result in canavanine-resistant (Can^R^) and adenine auxotroph clones (Ade-) that appear red when plated on selective media. The closest origin of replication is *ARS507*. As such, the lagging strand template for DNA replication is noted at the top of the figure. The CCTG and CAGG orientations refer to their placement on the lagging strand template. (B) Growth of strains on media containing canavanine and low adenine (5 μg/mL). (GAA)_5_ and (GAA)_220_ strains are described in REF. (C) CCTG/CAGG repeats elevate arm loss rates. Arm loss rates of (GAA)_5_, (CAGG)_100_, (CAGG)_138_, and (CCTG)100 from three-day incubation on selective media. Arm loss rates were calculated with FluCalc, which uses the Ma-Sandri-Sarkar maximum likelihood estimator model with a correction for sampling and plating efficiency. Error bars indicate standard error from three independent experiments. Statistical significance was evaluated by unpaired *t* test (* *p*<0.05).

We tested DNA fragility in (CCTG)_100_ and (CAGG)_100_ strains as well as a (CAGG)_138_ strain that was isolated as a spontaneous expansion during strain construction. We observed an increase in Can^R^Ade-clones that was above the background levels of a (GAA)_5_ negative control strain but noticeably less than (GAA)_220_ (Fig. 4b). Through fluctuation analyses, we quantified arm loss rates of the (CCTG)_100_ and (CAGG)_138_ orientations to be significantly increased compared to the (GAA)_5_ negative control (*p*<0.05) whereas (CAGG)_100_ showed an upward trend that was not significantly different (Fig. 4c). The rates of *CAN1* mutation alone among the CAGG/CCTG strains were not significantly different compared to the (GAA)_5_ negative control (Supplemental Fig. 6).

## Discussion

We have established an experimental system to characterize massive contractions of CCTG repeats, showing by genetic analysis that homologous recombination is required for this process. Through analysis of double mutants, we characterize epistatic relationships between genes involved in large-scale CCTG contractions. Furthermore, we provide the first evidence that CAGG/CCTG repeats increase DSB formation and chromosomal fragility *in vivo*.

Large-scale CCTG contractions were not dependent on proteins involved in lagging strand synthesis (Pol32) and Ozakaki fragment processing (Rad27) demonstrating that the genetic control is distinct from large-scale GAA repeat contractions, which were substantially increased in *pol32Δ* and *rad27Δ* mutants. However, we did see an increase in CCTG contractions in the Dna2-H547A mutant, which may involve Okazaki fragment processing by cleaving long 5’ flaps or another role in DSB processing (discussed below). GAA repeats form a triplex structure consisting of a GAA/TTC double helix with standard Watson Crick base pairing and, in its most stable configuration, a third GAA homopurine strand that anneals via Hoogsteen hydrogen bonds. Large contractions of GAA repeats were proposed through DNA polymerase occasionally bypassing this secondary structure (*32*). However, CAGG/CCTG repeats are not predicted to form triplex structures. Rather, there is gel electrophoretic evidence for hairpins (more stable in the CAGG orientation) (*21*) and slipped strand DNA (*22*). High resolution NMR studies with nucleotide substitution experiments have illuminated additional secondary structures such as dumbbells, minidumbbells (MDBs), and loops, which may interchange configurations rapidly. Though strand slippage on the template strand is likely to contribute to some repeat contractions, these are likely to be 1-3 unit deletion events.

Since we observe CCTG repeat contractions that exceed 80 repeat units, we propose a model for repeat contractions that is distinct from replication slippage (Fig. 5). Both camptothecin and hydroxyurea treatments elevate contraction rates even in the wild type strain. Thus, we propose that DSBs are generated during S phase DNA replication. Uncoupling of the replication fork helicase and polymerase could increase the persistence of ssDNA, secondary structure formation, and subsequent breaks. The DSBs could be generated by structure-specific nucleases acting at the distortions in B-DNA, though the nucleases remain to be identified. The DSBs are then repaired by homologous recombination, analogous to a fork restart mechanism. Because of the repetitive nature of the DNA template, out-of-register alignment will result in a massive contraction. Importantly, only events where the invading DNA end aligns towards the edge of the repeat tract on the template strand will result in a large enough repeat contraction for Ura+ selection. Next, we summarize some of our genetic analysis to describe how Rad51, Rad52, Sgs1, and Msh3 are required.

**Figure 5.**
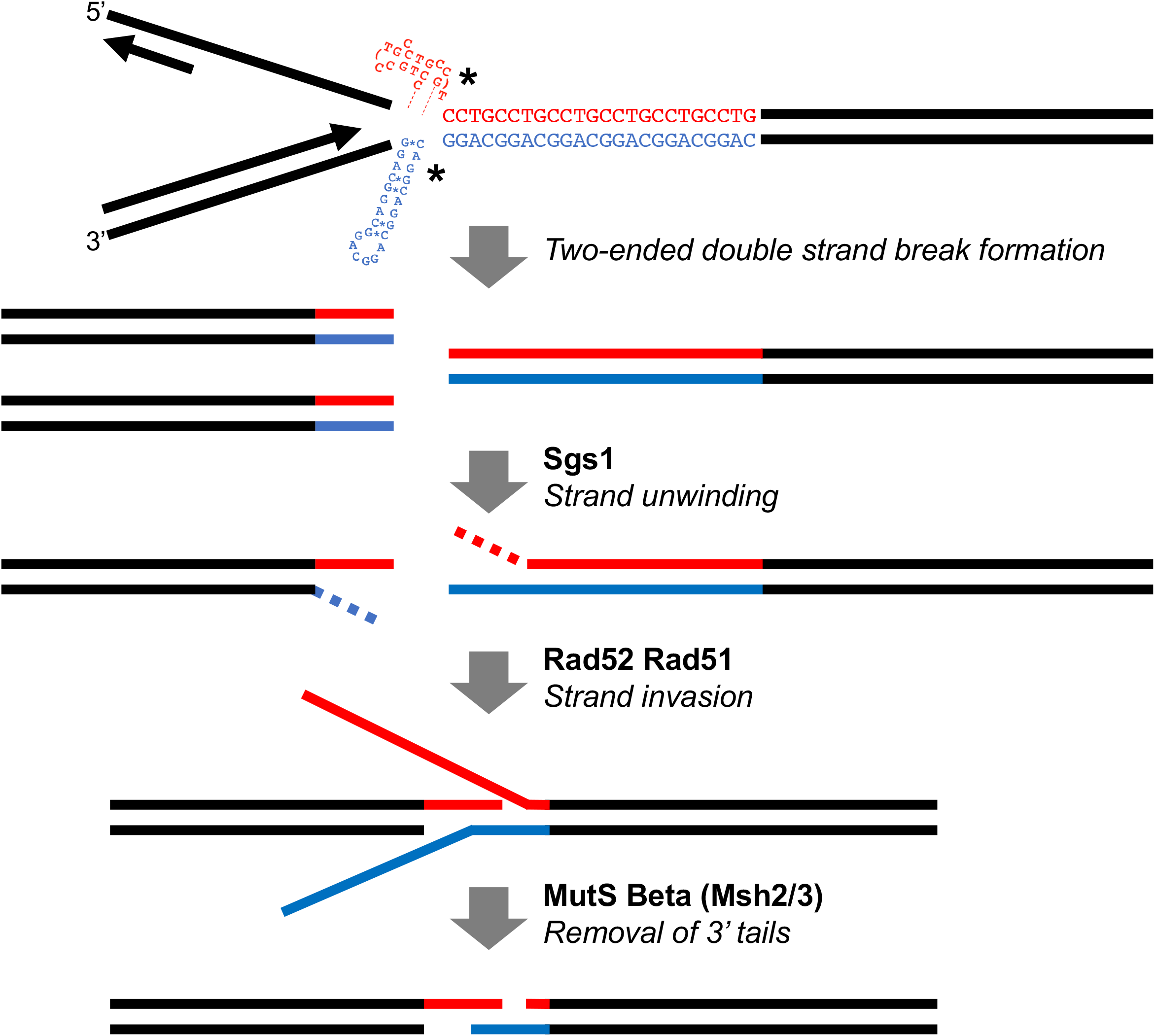
Proposed model of CCTG repeat contraction in budding yeast. During DNA replication, secondary structures such as dumbbells and hairpins form on single stranded DNA. Single stranded DNA formation could be elevated due to uncoupling of the replication fork at the DNA repeats (red/blue tracts). DNA double strand breaks occur at the CAGG/CCTG repeats, possibly mediated by structure-specific nucleases (asterisks). Sgs1 helicase promotes DNA strand unwinding to reveal 3’ single stranded DNA overhangs. Rad51 and Rad52 promote DNA invasion of the 3’ overhang to the homologous template. Because of the repetitive nature of the DNA template, out-of-register alignment will result in a massive contraction. Importantly, only events where the invading DNA end aligns towards the edge of the repeat tract on the template strand will result in a large enough repeat contraction for Ura+ selection. Following strand annealing, the resulting 3’ DNA tails are processed by MutS Beta.

Large-scale CCTG contractions were dependent on Rad51 and Rad52, though the effect was modest (2-fold decrease in knockouts). In a previous study examining single strand annealing within the ribosomal DNA array, *RAD52* was not essential, and it was proposed that the presence of large amounts of homology enable Rad52-independent DSB repair (*56*). The simple tandem repeat nature of the CCTG locus may make Rad51 and Rad52 dispensable for homology search leading to DSB repair. In our double mutant analysis, none of the effects were synergistic. Some double mutants were clearly epistatic (*rad51Δ rad52Δ* and *rad51Δ msh3Δ*) and others were epistatic or possibly additive (*rad51Δ sgs1Δ* and *sgs1Δ msh3Δ*). To harmonize these results, we interpret the genetic relationships with Sgs1 upstream of Rad51 and Rad51 upstream of Msh3.

Among its various roles in DNA repair, Sgs1 is known to promote 5’ DNA resection to form a longer 3’ single stranded DNA (ssDNA) tail (*45*). Sgs1 is also proposed to help translocation of the invading strand along a Holliday junction through interaction with Rad51, perhaps facilitating the search for homology. Because homology search at the CCTG repeats may not rely solely on Rad51 and Rad52, as described above, we investigated whether genes involved in DNA resection would also be required for large-scale CCTG contractions. We did not find an effect by deleting *MRE11* (MRX complex), *SAE2*, *RMI1* (STR complex), or *EXO1*. Thus, there may be redundant exonucleases that, along with Sgs1, generate the 3’ overhangs for homologous recombination. We tested the Dna2-H547A mutant that has been shown to have diminished endonuclease activity but unaffected ATPase and helicase activity (*47*). We found that contractions were increased 7.5-fold in the Dna2-H547A mutant, which was the opposite effect in the *sgs1Δ* mutant. The role of *DNA2*, which is essential, thus remains to be elucidated further. Because Sgs1 and Dna2 physically interact (*57*), one possibility is that Dna2 counteracts Sgs1 activity in promoting CCTG contractions.

MMR genes have been widely studied to investigate microsatellite DNA repeat instability, and their described roles depend on various factors including repeat sequence, length, and scale of expansion (*58*). We found that Msh3 is required for large-scale CCTG contractions, whereas Msh2 had no effect and Msh6 showed a slightly protective effect. As Msh2/3 forms a heterodimer, it is unclear why the *msh2Δ* mutant did not show a concomitant effect as *msh3Δ*. One possibility is that the Msh2/6 heterodimer plays an opposing role, which was proposed in a previous study investigating CAG/CTG repeat expansions. However, in that study, the absence of a phenotype in *msh2Δ* mutants was explained by opposing effects of Msh3 and Msh6 on lagging strand synthesis (*59*), a pathway we did not observe to play a major role in large-scale CCTG contractions. Msh2/3 has been shown to bind insertion/deletion mismatches in yeast (*60*). Thus, one possibility is that the heterodimer is involved in recognizing the DNA lesion that contributes to the DNA break. However, our double mutant analysis places *RAD51* genetically upstream of *MSH3*. Instead, we propose that Msh3 is involved in removal of the 3’ tail, like its described role in single strand annealing (*44, 61*). Characterizing the role of Msh3 warrants further investigation to clarify the role of MMR on large-scale CCTG contractions.

We have shown that disease-causing lengths of CCTG repeats cause chromosomal fragility *in vivo*. Because the assay requires DSB formation followed by DNA repair via telomere addition or break-induced replication with a non-homologous template, the rate of arm loss is likely to be an underestimate of total DSB formation induced by CCTG repeats. Given that CCTG repeat length can reach thousands in DM2 patients, it is intriguing to consider that DSB repair mechanisms might also contribute to such massive repeat expansions in human cells. Our system does not permit the selection of large-scale expansions since the starting strain is already Ura-. Thus, new experimental models will need to be developed to investigate CCTG repeat expansions in a systematic way. Conversely, it was shown that CCTG/CAGG tracts decreased significantly in the affected offspring of DM2 patients (*62*), and one intriguing possibility is that the contractions result from recombination following programmed DSBs during meiosis. One bioinformatics study proposes that the DM2 CCTG repeats may have originated from an AluSx element insertion (*63*). As long-read sequencing of repetitive DNA continues to advance, it will be intriguing to evaluate the prevalence of CCTG repeats in the human genome and their molecular properties of expansions, contractions, and DNA fragility.

## Acknowledgments

Research in the lab of J.C.K is supported by NIH SC3GM127198 with prior support from NIH K12GM074869. Undergraduate research support was provided by various institutions: DP and AD (CSUSM Summer Scholars), TH and SH (NIH T34 GM008807), BA (NSF REU 1852189), and MOO (NSF REU 1852189). LH was supported by a CSUPERB Graduate Student Research Restart award. We thank Kirill Lobachev for reagents, Alexandra Khristich for advice on mutant construction, and the Mirkin lab for helpful discussions. We thank Derrek Schartz for technical assistance during the preliminary stages of this project and Gaby Ramirez (NIH BRIDGES to Baccalaureate R25 GM066341) for help with plasmid cloning.

## Literature Cited

1. G. Meola, R. Cardani, Myotonic dystrophies: An update on clinical aspects, genetic, pathology, and molecular pathomechanisms. Biochimica et biophysica acta 1852, 594–606 (2015).

2. L. J. Sznajder, M. S. Swanson, Short Tandem Repeat Expansions and RNA-Mediated Pathogenesis in Myotonic Dystrophy. Int J Mol Sci 20, (2019).

3. T. Zu et al., Non-ATG-initiated translation directed by microsatellite expansions. Proc Natl Acad Sci U S A 108, 260–265 (2011).

4. T. Zu et al., RAN Translation Regulated by Muscleblind Proteins in Myotonic Dystrophy Type 2. Neuron 95, 1292–1305 e1295 (2017).

5. K. Usdin, N. C. House, C. H. Freudenreich, Repeat instability during DNA repair: Insights from model systems. Critical reviews in biochemistry and molecular biology 50, 142–167 (2015).

6. A. N. Khristich, S. M. Mirkin, On the wrong DNA track: Molecular mechanisms of repeat-mediated genome instability. The Journal of Biological Chemistry 295, 4134–4170 (2020).

7. A. A. Polyzos, C. T. McMurray, Close encounters: Moving along bumps, breaks, and bubbles on expanded trinucleotide tracts. DNA Repair (Amst) 56, 144–155 (2017).

8. V. C. Wheeler, V. Dion, Modifiers of CAG/CTG Repeat Instability: Insights from Mammalian Models. J Huntingtons Dis 10, 123–148 (2021).

9. C. E. Pearson, R. R. Sinden, Trinucleotide repeat DNA structures: dynamic mutations from dynamic DNA. Curr Opin Struct Biol 8, 321–330 (1998).

10. J. C. Kim, S. M. Mirkin, The balancing act of DNA repeat expansions. Current opinion in genetics & development 23, 280–288 (2013).

11. L. Poggi, G. F. Richard, Alternative DNA Structures In Vivo: Molecular Evidence and Remaining Questions. Microbiol Mol Biol Rev 85, (2021).

12. N. Fouche, S. Ozgur, D. Roy, J. D. Griffith, Replication fork regression in repetitive DNAs. Nucleic acids research 34, 6044–6050 (2006).

13. A. Kerrest et al., SRS2 and SGS1 prevent chromosomal breaks and stabilize triplet repeats by restraining recombination. Nature structural & molecular biology 16, 159–167 (2009).

14. J. L. Callahan, K. J. Andrews, V. A. Zakian, C. H. Freudenreich, Mutations in yeast replication proteins that increase CAG/CTG expansions also increase repeat fragility. Mol Cell Biol 23, 7849–7860 (2003).

15. E. J. Polleys, C. H. Freudenreich, Genetic Assays to Study Repeat Fragility in Saccharomyces cerevisiae. Methods Mol Biol 2056, 83–101 (2020).

16. E. J. Polleys, C. H. Freudenreich, Homologous recombination within repetitive DNA. Current opinion in genetics & development 71, 143–153 (2021).

17. M. Izzo et al., Molecular Therapies for Myotonic Dystrophy Type 1: From Small Drugs to Gene Editing. Int J Mol Sci 23, (2022).

18. M. Nakamori et al., A slipped-CAG DNA-binding small molecule induces trinucleotide-repeat contractions in vivo. Nat Genet 52, 146–159 (2020).

19. C. Cinesi, L. Aeschbach, B. Yang, V. Dion, Contracting CAG/CTG repeats using the CRISPR-Cas9 nickase. Nat Commun 7, 13272 (2016).

20. C. L. Liquori et al., Myotonic dystrophy type 2 caused by a CCTG expansion in intron 1 of ZNF9. Science 293, 864–867 (2001).

21. R. Dere, M. Napierala, L. P. Ranum, R. D. Wells, Hairpin structure-forming propensity of the (CCTG.CAGG) tetranucleotide repeats contributes to the genetic instability associated with myotonic dystrophy type 2. The Journal of Biological Chemistry 279, 41715–41726 (2004).

22. S. F. Edwards, M. Sirito, R. Krahe, R. R. Sinden, A Z-DNA sequence reduces slipped-strand structure formation in the myotonic dystrophy type 2 (CCTG) x (CAGG) repeat. Proc Natl Acad Sci U S A 106, 3270–3275 (2009).

23. S. L. Lam, F. Wu, H. Yang, L. M. Chi, The origin of genetic instability in CCTG repeats. Nucleic acids research 39, 6260–6268 (2011).

24. P. Guo, S. L. Lam, Minidumbbell: A New Form of Native DNA Structure. Journal of the American Chemical Society 138, 12534–12540 (2016).

25. P. Guo, S. L. Lam, Unusual structures of CCTG repeats and their participation in repeat expansion. Biomolecular concepts 7, 331–340 (2016).

26. R. Dere, R. D. Wells, DM2 CCTG*CAGG repeats are crossover hotspots that are more prone to expansions than the DM1 CTG*CAG repeats in Escherichia coli. J Mol Biol 360, 21–36 (2006).

27. S. F. Edwards, V. I. Hashem, E. A. Klysik, R. R. Sinden, Genetic instabilities of (CCTG).(CAGG) and (ATTCT).(AGAAT) disease-associated repeats reveal multiple pathways for repeat deletion. Molecular carcinogenesis 48, 336–349 (2009).

28. M. M. Krasilnikova, S. M. Mirkin, Replication stalling at Friedreich’s ataxia (GAA)n repeats in vivo. Mol Cell Biol 24, 2286–2295 (2004).

29. E. Grabczyk, K. Usdin, Generation of microgram quantities of trinucleotide repeat tracts of defined length, interspersion pattern, and orientation. Anal Biochem 267, 241–243 (1999).

30. K. A. Shah et al., Role of DNA polymerases in repeat-mediated genome instability. Cell reports 2, 1088–1095 (2012).

31. A. A. Shishkin et al., Large-scale expansions of Friedreich’s ataxia GAA repeats in yeast. Molecular cell 35, 82–92 (2009).

32. A. N. Khristich, J. F. Armenia, R. M. Matera, A. A. Kolchinski, S. M. Mirkin, Large-scale contractions of Friedreich’s ataxia GAA repeats in yeast occur during DNA replication due to their triplex-forming ability. Proc Natl Acad Sci U S A 117, 1628–1637 (2020).

33. M. Napierala, P. Parniewski, A. Pluciennik, R. D. Wells, Long CTG.CAG repeat sequences markedly stimulate intramolecular recombination. The Journal of Biological Chemistry 277, 34087–34100 (2002).

34. M. Napierala, R. Dere, A. Vetcher, R. D. Wells, Structure-dependent recombination hot spot activity of GAA.TTC sequences from intron 1 of the Friedreich’s ataxia gene. The Journal of Biological Chemistry 279, 6444–6454 (2004).

35. J. J. Miret, L. Pessoa-Brandao, R. S. Lahue, Orientation-dependent and sequence-specific expansions of CTG/CAG trinucleotide repeats in Saccharomyces cerevisiae. Proc Natl Acad Sci U S A 95, 12438–12443 (1998).

36. S. Bhattacharyya, R. S. Lahue, Saccharomyces cerevisiae Srs2 DNA helicase selectively blocks expansions of trinucleotide repeats. Molecular and Cellular Biology 24, 7324–7330 (2004).

37. R. Sundararajan, L. Gellon, R. M. Zunder, C. H. Freudenreich, Double-strand break repair pathways protect against CAG/CTG repeat expansions, contractions and repeat-mediated chromosomal fragility in Saccharomyces cerevisiae. Genetics 184, 65–77 (2010).

38. X. A. Su, V. Dion, S. M. Gasser, C. H. Freudenreich, Regulation of recombination at yeast nuclear pores controls repair and triplet repeat stability. Genes Dev 29, 1006–1017 (2015).

39. J. C. Kim, S. T. Harris, T. Dinter, K. A. Shah, S. M. Mirkin, The role of break-induced replication in large-scale expansions of (CAG)n/(CTG)n repeats. Nature structural & molecular biology 24, 55–60 (2017).

40. C. Seong, S. Colavito, Y. Kwon, P. Sung, L. Krejci, Regulation of Rad51 recombinase presynaptic filament assembly via interactions with the Rad52 mediator and the Srs2 anti-recombinase. The Journal of Biological Chemistry 284, 24363–24371 (2009).

41. S. Bhattacharyya, R. S. Lahue, Srs2 helicase of Saccharomyces cerevisiae selectively unwinds triplet repeat DNA. The Journal of Biological Chemistry 280, 33311–33317 (2005).

42. A. Kerrest et al., SRS2 and SGS1 prevent chromosomal breaks and stabilize triplet repeats by restraining recombination. Nat Struct Molec Biol 16, 159–167 (2009).

43. S. V. Gupta, K. H. Schmidt, Maintenance of Yeast Genome Integrity by RecQ Family DNA Helicases. Genes (Basel) 11, (2020).

44. N. Sugawara, T. Goldfarb, B. Studamire, E. Alani, J. E. Haber, Heteroduplex rejection during single-strand annealing requires Sgs1 helicase and mismatch repair proteins Msh2 and Msh6 but not Pms1. Proc Natl Acad Sci U S A 101, 9315–9320 (2004).

45. P. Cejka, S. C. Kowalczykowski, The full-length Saccharomyces cerevisiae Sgs1 protein is a vigorous DNA helicase that preferentially unwinds holliday junctions. The Journal of Biological Chemistry 285, 8290–8301 (2010).

46. M. D. Huber, D. C. Lee, N. Maizels, G4 DNA unwinding by BLM and Sgs1p: substrate specificity and substrate-specific inhibition. Nucleic acids research 30, 3954–3961 (2002).

47. K. H. Lee et al., The endonuclease activity of the yeast Dna2 enzyme is essential in vivo. Nucleic acids research 28, 2873–2881 (2000).

48. G. M. Williams, J. A. Surtees, MSH3 Promotes Dynamic Behavior of Trinucleotide Repeat Tracts In Vivo. Genetics 200, 737–754 (2015).

49. K. Manley, T. L. Shirley, L. Flaherty, A. Messer, Msh2 deficiency prevents in vivo somatic instability of the CAG repeat in Huntington disease transgenic mice. Nat Genet 23, 471–473 (1999).

50. W. J. van den Broek et al., Somatic expansion behaviour of the (CTG)n repeat in myotonic dystrophy knock-in mice is differentially affected by Msh3 and Msh6 mismatch-repair proteins. Human molecular genetics 11, 191–198 (2002).

51. L. Foiry et al., Msh3 is a limiting factor in the formation of intergenerational CTG expansions in DM1 transgenic mice. Hum Genet 119, 520–526 (2006).

52. A. M. Gannon, A. Frizzell, E. Healy, R. S. Lahue, MutSbeta and histone deacetylase complexes promote expansions of trinucleotide repeats in human cells. Nucleic acids research 40, 10324–10333 (2012).

53. G. Pannafino, E. Alani, Coordinated and Independent Roles for MLH Subunits in DNA Repair. Cells 10, (2021).

54. H. M. Kim et al., Chromosome fragility at GAA tracts in yeast depends on repeat orientation and requires mismatch repair. The EMBO journal 27, 2896–2906 (2008).

55. Y. Zhang, N. Saini, Z. Sheng, K. S. Lobachev, Genome-wide screen reveals replication pathway for quasi-palindrome fragility dependent on homologous recombination. PLoS genetics 9, e1003979 (2013).

56. B. A. Ozenberger, G. S. Roeder, A unique pathway of double-strand break repair operates in tandemly repeated genes. Mol Cell Biol 11, 1222–1231 (1991).

57. P. Cejka et al., DNA end resection by Dna2-Sgs1-RPA and its stimulation by Top3-Rmi1 and Mre11-Rad50-Xrs2. Nature 467, 112–116 (2010).

58. E. A. Sia, R. J. Kokoska, M. Dominska, P. Greenwell, T. D. Petes, Microsatellite instability in yeast: dependence on repeat unit size and DNA mismatch repair genes. Mol Cell Biol 17, 2851–2858 (1997).

59. A. Kantartzis et al., Msh2-Msh3 interferes with Okazaki fragment processing to promote trinucleotide repeat expansions. Cell reports 2, 216–222 (2012).

60. Y. Habraken, P. Sung, L. Prakash, S. Prakash, Binding of insertion/deletion DNA mismatches by the heterodimer of yeast mismatch repair proteins MSH2 and MSH3. Curr Biol 6, 1185–1187 (1996).

61. R. Bhargava, D. O. Onyango, J. M. Stark, Regulation of Single-Strand Annealing and its Role in Genome Maintenance. Trends Genet 32, 566–575 (2016).

62. J. W. Day, et al., Myotonic dystrophy type 2: molecular, diagnostic and clinical spectrum. Neurology 60, 657–664 (2003).

63. T. Kurosaki et al., The unstable CCTG repeat responsible for myotonic dystrophy type 2 originates from an AluSx element insertion into an early primate genome. PLoS One 7, e38379 (2012).

